# Heterogenous mitochondrial ultrastructure and metabolism of human glioblastoma cells: differences between stem-like and differentiated cancer cells in response to chemotherapy

**DOI:** 10.1101/2025.02.17.638633

**Authors:** Urban Bogataj, Metka Novak, Simona Katrin Galun, Klementina Fon Tacer, Miloš Vittori, Cornelis J.F. Van Noorden, Barbara Breznik

## Abstract

**Background:** Glioblastoma stem-like cells (GSCs) contribute to the resistance of glioblastoma (GBM) tumors to standard therapies. The cellular and molecular background of the resistance of GSCs to the chemotherapeutic agent temozolomide is not yet fully understood, in particular in the context of cellular metabolism and the role of mitochondria. The aim of this study was to perform a detailed ultrastructural characterization of the mitochondria of GSCs prior and post temozolomide exposure and to compare it to differentiated GBM cells.

**Methods:** Patient-derived and established GSC and GBM differentiated cell lines were used for the study. The ultrastructure of the mitochondria of the examined cell lines was assessed by transmission electron microscopy. The microscopic analysis was complemented and compared by an analysis of cell metabolism using cell viability assay and extracellular flux analysis using Seahorse assay.

**Results:** We found that the metabolic profile of GSCs is quiescent and aerobic. Their elongated mitochondria with highly organized cristae is indicating increased biogenesis and mitochondrial fusion and corresponds to a more OXPHOS-dependent metabolism. The metabolism of GSCs is dependent on OXPHOS and there are no changes in defective mitochondria fraction after the treatment with temozolomide. In contrast, differentiated GBM cells with fragmented mitochondria, which have less organized cristae, are more energetic and glycolytic. Temozolomide treatment induced significant ultrastructural mitochondrial damage in differentiated GBM cells and had less effect on mitochondria in GSCs, suggesting that mitochondria play an important role in the resistance of GSCs to temozolomide treatment.

**Conclusions:** We demonstrated differences in mitochondrial ultrastructure and cellular metabolism between GSCs and differentiated GBM cells in response to temozolomide. This study provides a basis for further studies addressing the chemotherapy resistance of GSCs and the effects of different treatment regimens on the mitochondrial structure and function of GSCs.

**Graphical Abstract:** 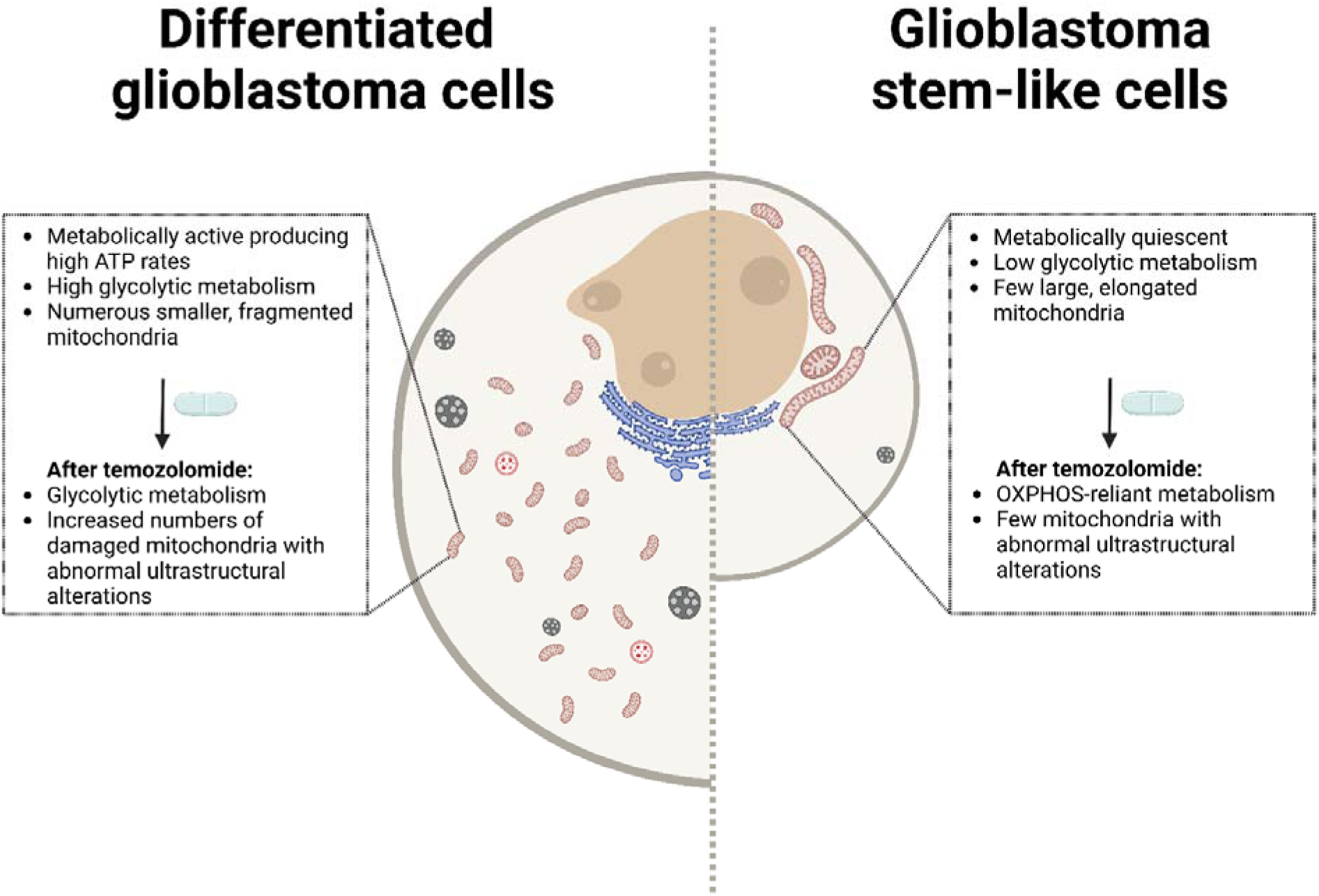

## Introduction

Glioblastoma (GBM), the most common primary brain tumor in adults, remains one of the most aggressive malignancies with median survival of 16 months after diagnosis [1] and so far, incurable. Standard treatment includes maximal safe surgical removal of tumor tissue, radiotherapy and chemotherapy using the alkylating drug temozolomide (TMZ) [2]. However, due to GBM heterogeneity, its invasive nature, and its resistance to chemotherapy and radiotherapy, the GBM almost always reoccurs in a more aggressive form [1].

An important general characteristic of GBM is that it involves numerous heterogenous cell types including the glioma stem-like cells (GSCs) [3, 4] that exhibit diverse metabolic profiles [5, 6]. A consequence is enormous plasticity and adaptability to therapeutic interventions resulting in recurrence and resistance to therapy. GSCs play a crucial role in therapy resistance [6]. GSCs reside within specific niches within tumors that are primarily hypoxic [4, 7–9]. The hypoxic tumor microenvironment is important for the maintenance of GSCs [10, 11] but requires specific metabolic adaptations, with mitochondria playing a crucial role in the maintenance of GSCs [12].

One of the common characteristics of cancer cell metabolism is the Warburg effect, which designates that cancer cells often rely primarily on glycolysis to produce ATP even in the presence of oxygen. Dysfunctional mitochondria in cancer cells were suggested as one of the major culprits [13–15], however, recent data suggest that cancer cells and, in particularly cancer stem cells [16], including GSCs [5, 17], often rather rely on oxidative phosphorylation (OXPHOS) to produce ATP [18–20]. The mitochondria are thus crucial in cancer cell biology through their direct involvement in OXPHOS, besides other roles they play in cell signaling, synthesis of macromolecules, oxidative stress and apoptosis [21–23]. Mitochondrial fusion and elongated mitochondria support efficient OXPHOS especially during nutrient withdrawal, whereas fragmented mitochondria are often associated with impaired OXPHOS and nutrient excess [24]. Cancer cells generally exhibit fragmented mitochondria, associated with low OXPHOS activity and increased glycolysis [25–27]. Cancer stem cells, however, exhibit increased biogenesis of mitochondria, consistent with increased OXPHOS [28, 29], however, their role in therapy response and resistance is not well understood.

The ultrastructure of mitochondria in GBM cells is very variable, but a consistent observation is the presence of abundant swollen and electron-lucent mitochondria that display reduced and disorganized cristae [18, 19, 30, 31]. So far, most ultrastructural analysis of GBM cells have been performed in tumor biopsies [32, 33] and tissue explants obtained from patients during tumor excision [34]. These studies predominately focus on the cells that form the bulk of tumor mass and do not specifically address GSC properties and their response to chemotherapy.

Given the importance of GSC in therapy resistance, the aim of our study was to provide a detailed ultrastructural characterization of cultured GSCs upon exposure to the chemotherapeutic TMZ and compare it to differentiated GBM cells. The microscopic analysis was complemented and compared with cellular metabolism analysis using cell viability assay and extracellular flux analysis. A commonly used method for measuring cellular metabolism is extracellular flux (XF; also known as Seahorse) analysis. XF analysis measures extracellular acidification (ECAR) and oxygen consumption (OCR) rates as markers of glycolysis and mitochondrial OXPHOS, respectively [35, 36], and was applied to evaluate cellular metabolism in GSCs and differentiated GBM cells.

We provide novel insights into structural and metabolic responses of GBM and GSC cells to chemotherapy and lay foundation for further studies addressing GSC therapeutic resistance and the effects of different treatment regimens on the mitochondrial structure and function of GSCs.

## Materials and methods

### Cell cultures

In this study, two GSC lines (NCH644 and NCH421k) and two differentiated GBM cell cultures (U87 and NIB140) were included. Commercially available GSCs NCH421k and NCH644 were purchased from the Cell Lines Service (CLS) GmbH, Eppelheim, Germany. Cells were grown in serum free conditions in Neurobasal Medium as described before [37]. Cell line U87 MG was obtained from the American Type Culture Collection (ATCC, Manassas, VA, USA) and was cultured in low glucose DMEM supplemented with 10% fetal bovine serum (FBS; Gibco, Thermo Fisher Scientific, Waltham, MA, USA), 2 mM L-glutamine and 1% penicillin/streptomycin (both: Sigma Aldrich, St. Louis, MO, USA). NIB140 are patient-derived cell cultures from fresh GBM tissues and were established as described before [38–40]. NIB140 cells were grown in Dulbecco’s modified Eagle’s medium (DMEM) (Hyclone, GE Healthcare, Chicago, IL, USA) supplemented with 10% FBS (Gibco), 2 mM L-glutamine, and 1× penicillin/streptomycin (both: Sigma Aldrich). Growing cells were detached with a 0.25% trypsin EDTA solution (Gibco). NIB140 cells express GBM cell markers [39, 41]. All cells were cultured at 37 °C, in the presence of 5% CO_2_ and 95% humidity and checked for Mycoplasma using MycoAlert Mycoplasma Detection Kit (Lonza, Basel, Switzerland). Authentication of cells was performed by DNA fingerprinting using Amp-FlSTR Profiler Plus PCR Amplification Kit, as described previously [42].

### Cell viability assay

Viability of cells was determined after 48 h of treatment with TMZ (Sigma-Aldrich) using the MTT (3-(4,5-dimethylthiazol-2-yl)-2,5-diphenyltetrazolium-bromide; Sigma-Aldrich) reagent for U87 and NIB140 cells and MTS (3-(4,5-dimethylthiazol-2-yl)-5-(3-carboxymethoxyphenyl)-2-(4-sulfophenyl)-2H-tetrazolium; Promega, Madison, WI, USA) reagent for NCH cells. Assays were performed according to the manufacturer’s instructions. Briefly, cells were seeded into 96-well plates (8000 cells/well) and grown overnight. Cells were treated with different concentrations of TMZ (25–400 µM). Stock solutions of TMZ were prepared in dimethyl sulfoxide (DMSO, Sigma-Aldrich). Control incubation media contained the same amount of vehicle DMSO (0.9%, v/v). After 48 h, MTT or MTS was added and 3 h after incubation at, absorbance was measured as the change in optical density (ΔOD 570/690 nm) using a microplate reader (Synergy™ HT, Bio Tek Instruments Inc., Winooski, VT, USA). Cell viability data were analyzed using GraphPad Prism software (GraphPad Software, San Diego, CA, USA) and presented as % of vehicle control.

### Extracellular flux analysis

Extracellular flux (XF) analysis was performed using Seahorse XFe24 Flux Analyzer (Agilent Technologies, Santa Clara, CA, USA) to measure the oxygen consumption rate (OCR) and extracellular acidification rate (ECAR) of live cells in a 24-well plate format. GBM cells were grown in T25 flask as controls (0.5% DMSO) or treated with 100 µM of TMZ for 48 h. Before the XF experiment, cells were harvested, 80,000 cells/well were seeded in poly-d-Lysine (Poly-D-lysine hydrobromide, Sigma)-coated XF24 cell culture microplates in Seahorse XF DMEM medium (pH = 7.4 with 10 mM XF glucose, 1 mM XF pyruvate and 2 mM XF glutamine) in duplicates and centrifuged to allow the cells to attach to the bottom of the plates. The cells were transferred to a CO_2_-free incubator at 37 °C for 20 min. During this time, assays were prepared, and cartridges loaded. XF Real-time ATP Rate Assay Kit (#103592; Agilent), XF Glycolytic Rate Assay Kit (#103344; Agilent) and XF Cell Mito Stress Test Kit (#103015; Agilent) were applied according to manufacturer’s instructions. Immediately after the run, the cells were fixed with 4% paraformaldehyde (PFA; Sigma) and incubated with Hoechst 33442 (Sigma) to count the number of cells in each well for normalization of the XF results. Hoechst signal was measured using Cytation 5 Cell Imaging multi-mode reader (Bio Tek) and cell counts were analyzed using Gen 5 software (Bio Tek). Flux rates were normalized to cell counts in each well. Data were analyzed using Wave software version 2.6.1 and an online software version at the Agilent cloud (https://seahorseanalytics.agilent.com/). Statistically significant differences in metabolic rates and parameters between different cells and between control and TMZ-treated samples were analyzed using GraphPad Prism software (GraphPad Software).

### Transmission electron microscopy

#### Sample preparation

The cells were exposed either to 100 μM TMZ (Sigma-Aldrich) or 0.9%, v/v DMSO (Sigma-Aldrich) (vehicle) for 48 h. Then, the cells were fixed using 2.5% glutaraldehyde (SPI, West Chester, PA, USA) and 2% PFA (Sigma-Aldrich) in PBS (Gibco). Following fixation, the cells were rinsed in PBS and postfixed in 1% OsO_4_ (SPI) in PBS (Gibco). Then, the cells were washed in PBS (Gibco) and embedded in 2% low melting point agarose (Sigma-Aldrich). The blocks of agarose with embedded cells were solidified on ice and cut into small pieces (<1 mm in smallest dimension). The pieces of agarose with cells were then dehydrated in graded series of ethanol (Honeywell, Seelze, Germany) and acetone (Merck, Darmstadt, Germany). Finally, the samples were embedded in epoxy resin Agar 100 (Agar Scientific, Rotherham, UK). The resin was polymerized for 24 h at 60 °C in embedding molds. Semithin and ultrathin sections were prepared with a Reichert Ultracut S (Leica, Wetzlar, Germany) ultramicrotome, using glass and diamond knives. The semithin sections were stained with Azure II (Merck) – methylene blue (Sigma-Aldrich) and examined with a light microscope. The ultrathin sections were contrasted with uranyl acetate (SPI) and lead citrate (SPI).

#### Imaging

Contrasted ultrathin sections were imaged with a CM100 transmission electron microscope (Philips, Eindhoven, The Netherlands). The electron micrographs were acquired with an Orius 200 camera (Gatan, Pleasanton, CA, USA) and Digital Micrograph software (Gatan). Five cells were imaged for each biological replicate per cell line. Numerous partly overlapping images were acquired per cell at 3 different magnifications. The acquired images were stitched together with FIJI [43] open-source software and TrakEM2 [44] plugin to obtain images with large field of view and high resolution. Stitched images at lowest magnification covered whole cells at sufficient resolution to obtain a general overview of cell ultrastructure and to perform stereological quantification of mitochondrial and nuclear volume densities. Stitched images at medium magnification covered large areas of cytoplasm with sufficient resolution to obtain overall ultrastructural features of individual cellular organelles and to perform measurements of mitochondrial length and width. Stitched images at highest magnification covered small areas of cytoplasm with sufficient resolution to distinguish fine ultrastructural features of mitochondria, such as the morphology of cristae.

#### Image analysis

We counted the number of mitochondrial cross-sections on stitched images covering whole cells using FIJI software with Cell counter plugin. For the stereological quantification of mitochondrial and nuclear volume density, an overlay of multipurpose grid of test points was made on stitched images covering whole analyzed cells. We used the multipurpose grid that is available as a macro for ImageJ/FIJI at: https://imagej.nih.gov/ij/macros/Multipurpose_grid.txt. In the test grid parameters, we set the density of test points to 1 point per 0.5 µm^2^. We counted the number of test points inside mitochondria, cell nuclei and other areas of cytoplasm with aid of the Cell counter plugin. From the obtained counts, we calculated the estimates for the volume density (volume fraction) of mitochondria and the volume density of cell nuclei according to the equation: 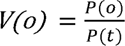, where the *V(o)* is the volume density of organelle in question, *P(o)* is the count of test points inside the organelles in question and the *P(t)* is the total count of points inside cells (Weibel et al. 1966). To describe the general morphology of the mitochondrial network analyzed in GBM cell lines, we measured the average length and width of mitochondrial cross-sections in stitched images at medium magnification. Then, we calculated the length:width ratio of mitochondrial cross-sections. If this ratio was close to 1, it indicated that the mitochondrial network consisted primarily of round mitochondria and was likely more fragmented. If this ratio was much greater than 1, it indicated that the mitochondrial network consisted of elongated mitochondria in a more interconnected mitochondrial network.

To investigate the effect of TMZ treatment on mitochondrial ultrastructure, features of mitochondria in 5 cells per biological replicate were evaluated. Based on their ultrastructural features, we classified the mitochondria into two following types: normal and defective. Normal mitochondria are defined as mitochondria with electron-dense matrix with narrow cristae (a) and mitochondria with electron-dense matrix with dilated cristae (b). Defective mitochondria are defined as mitochondria with electron-lucent matrix and cristae reduced in size and numbers (c) and mitochondria with abnormal ultrastructural alterations (swollen and containing membrane swirls) (d). The percentages/fractions of mitochondrial types in each cell were calculated from the total fraction of mitochondria with clearly visible ultrastructure at a 3400x magnification.

### MitoTracker staining

U87 and NIB140 cells were cultured on poly-L-lysine-coated coverslips in 24-well plates (Corning, NY, USA) and NCH421k cells were dissociated before stained using MitoTracker® Orange CMTMRos (Invitrogen, Thermo Fisher Scientific) according to the manufacturer’s instructions. Stained cells were fixed in 4% PFA in PBS. Following fixation, the cells were rinsed in PBS and stained with Hoechst 33442 (Sigma) to visualize cell nuclei. Imaging was performed with an AxioImager Z.1 microscope equipped with an AxioCam MRm camera using Axiovision software (all from Zeiss, Oberkochen, Germany).

### Statistical analysis

To compare morphometric parameters between the analyzed GBM cell lines quantitative measurements in 5 biological replicates (n = 5) of each cell line were performed. For each biological replicate we analyzed 5 cells. The following morphometric parameters were investigated: the number of mitochondrial cross-sections, mitochondrial volume density, nuclear volume density and length/width ratio of mitochondria. The statistical analyses were performed in the R statistical software environment. To test for statistically significant differences between cell lines, the analysis of variance (ANOVA) followed by Tukey’s test for pairwise comparisons was performed. Before the ANOVA analysis, we prepared Q-Q plots of residuals and performed a Shapiro-Wilk test to check the normality of data. We also performed Levene’s test to check the equality of variances.

To compare fractions of different mitochondrial types and volume density of mitochondria between TMZ-treated and non-treated cells, the experiments were performed in two independent repeats (n = 2). The statistical analyses were performed in GraphPad Prism software. To test for statistically significant differences in fractions of mitochondrial types, a Fisher’s exact test was performed. To test for statistically significant differences in volume density of mitochondria, an unpaired t-test was performed.

To compare metabolic profiles of GBM cells and GSCs, Seahorse analysis experiments were performed in 3 independent repeats (n = 3) and the statistical analyses were performed in GraphPad Prism software. Statistics were performed using Tukey’s multiple comparisons test. Differences in ATP production rates between TMZ-treated and non-treated samples within each cell line were statistically evaluated using an unpaired t-test. Tukey’s multiple comparisons test was used to compare all conditions after treatment with TMZ.

## Results

### Overall cellular ultrastructure

Both types of GSCs, NCH421k and NCH644, consisted of small cells of relatively homogeneous size and shape that aggregated in culture in small clusters. Both types of differentiated GBM cells, NIB140 and U87, were highly heterogeneous in terms of size and shape. Our stereological quantification of cell nuclear volume density indicated that the cell nuclei in GSCs occupied a significantly larger portion of cell volume than nuclei in differentiated GBM cells (Figure **1A**). GSCs had oval cell nuclei with abundant heterochromatin and a single nucleolus (Supplementary Figures **S1A, B**). In differentiated GBM cells the cell nuclei contained hardly any heterochromatin and usually multiple nucleoli (Supplementary Figures **S1C, D**). We observed that the cisternae of the endoplasmic reticulum and Golgi stacks were considerably more abundant in differentiated GBM cells than in GSCs. The differentiated GBM cells also contained abundant multivesicular bodies and secondary lysosomes, which were scarce in GSCs (Supplementary Figure **S2**).

**Figure 1:**
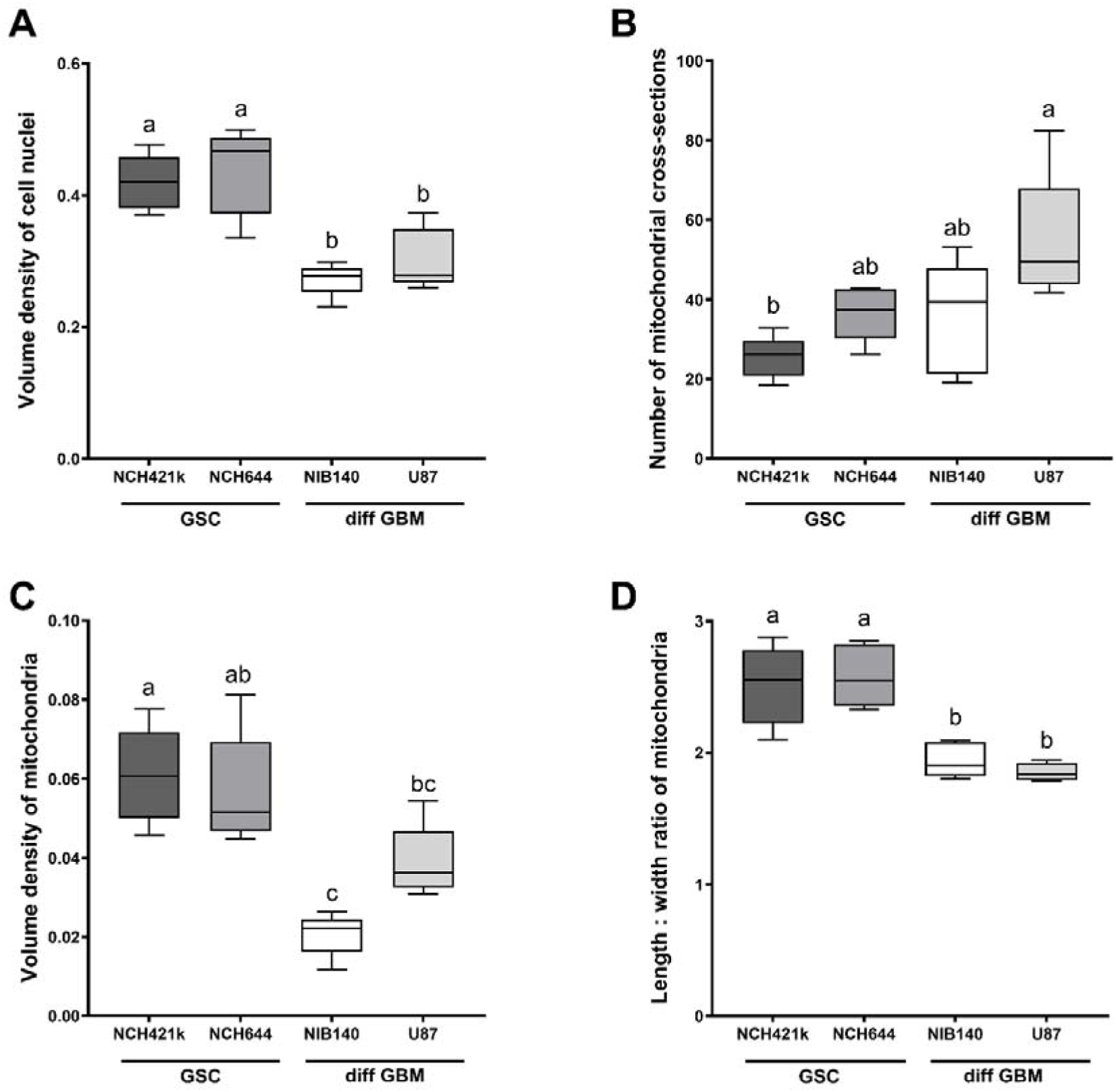
Quantification of **A**) volume density of cell nuclei, **B**) number of mitochondrial cross-sections per cell, **C**) volume density of mitochondria and **D**) length/width ratio of mitochondrial cross-sections. Data are represented by mean values ± SEM. Statistically significant differences are displayed with compact letters display (a, b, c). Cell lines that do not share any common letter have statistically significantly different means of a dependent variable (p < 0.05). Cell lines that share a common letter do not have statistically significant different means of a dependent variable (p ≥ 0.05).

### Mitochondrial ultrastructure differs between GBM cells

Quantification of mitochondrial cross-sections indicated a significant difference in mitochondrial numbers per cell between GSCs NCH421k and differentiated GBM cells U87 (Figure 1B), with mitochondria being more numerous in the U87 cells than in the NCH421k GSCs. Stereological quantification of mitochondrial volume density showed that the mitochondria in the differentiated GBM cells NIB140 occupied a significantly smaller proportion of cell cytoplasm than in both GSC lines (Figure 1C). Mitochondria in the differentiated GBM cells U87 occupied a significantly smaller proportion of cell cytoplasm than in the GSCs NCH421k. The quantification of ratio between mitochondrial length and width showed that the mitochondria in GSCs were more elongated than in differentiated GBM cells (Figure 1D).

The GSCs contained in general more large mitochondria and less smaller mitochondria, whereas differentiated GBM cells contained larger numbers of smaller mitochondria. In GSCs NCH644 and NCH421k, the most prevalent morphological type of mitochondria was represented by elongated, electron-dense mitochondria with narrow cristae which were oriented along the longitudinal axis of mitochondria (Figures **2A, C**). Occasionally we observed dilated cristae (Figure **2B**). In both types of GSCs, mitochondria occasionally displayed deformed cristae with swirled shape (Figure **2D**). In differentiated GBM cells U87 and NIB140, the mitochondria were rounder than in GSC lines and cristae were not organized, but rather oriented in variable directions (Figure **3**). The two most prevalent morphological types of mitochondria in differentiated GBM cells were electron-lucent swollen mitochondria with reduced numbers of narrow cristae (Figures **3A, C**) and electron-dense mitochondria with numerous dilated cristae (Figures **3B, D**). In U87 cells, we frequently observed electron-dense mitochondria with narrow cristae. MitoTracker staining confirmed dense active mitochondria in the cells (Supplementary Figure **S3**).

**Figure 2:**
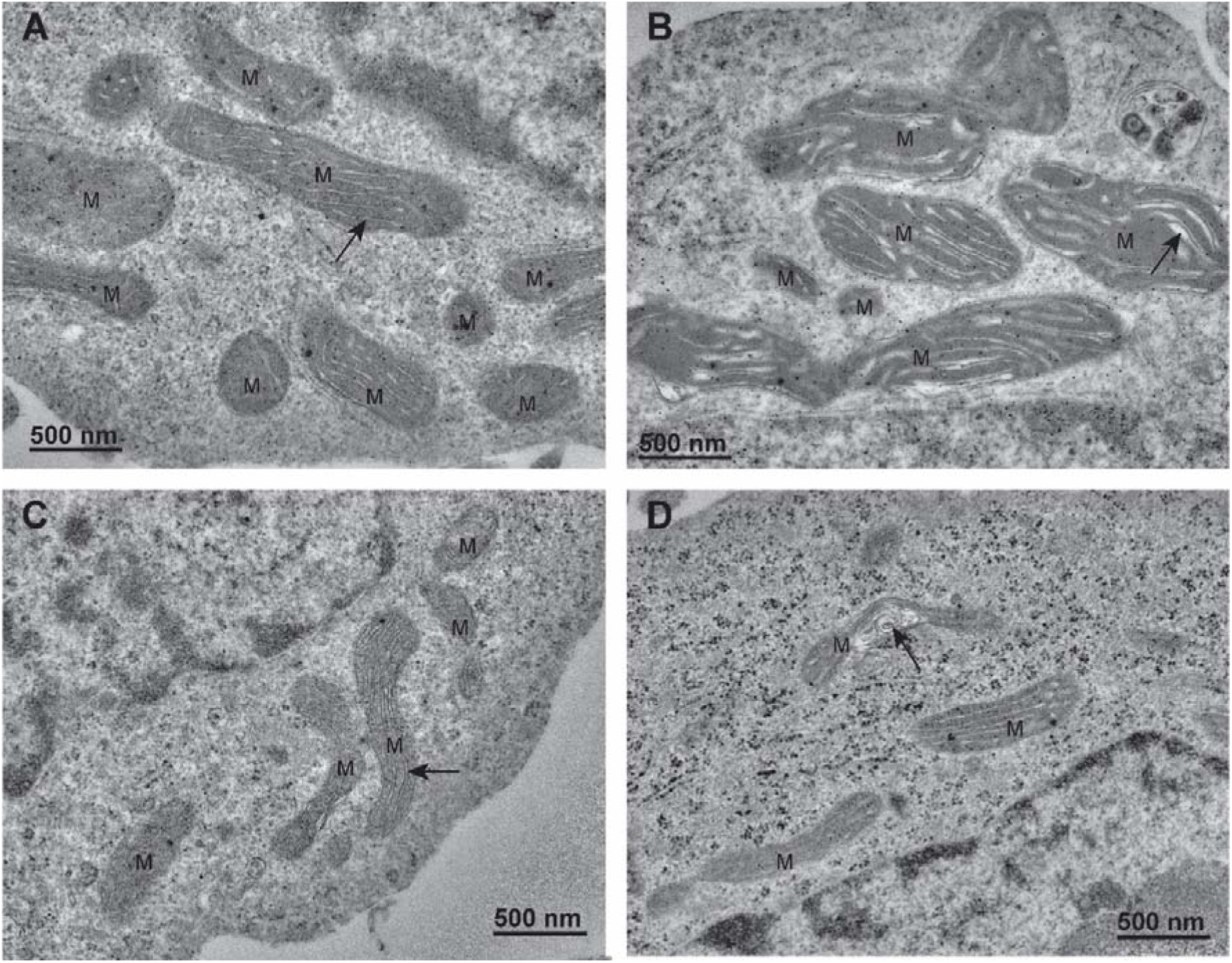
Mitochondria in GSCs have elongated mitochondria. **A**) Electron-dense mitochondria (M) with narrow cristae (arrow) oriented along the longitudinal axis of mitochondria in the NCH421k GSCs. **B**) Electron-dense mitochondria (M) with dilated cristae (arrow) in the NCH421k GSCs. **C**) Typical electron-dense mitochondria (M) with narrow cristae (arrow) oriented along the longitudinal axis of mitochondria in the NCH644 GSCs. **D**) Deformed cristae with swirled shape (arrow) in a mitochondrion of a NCH644 cells.

**Figure 3:**
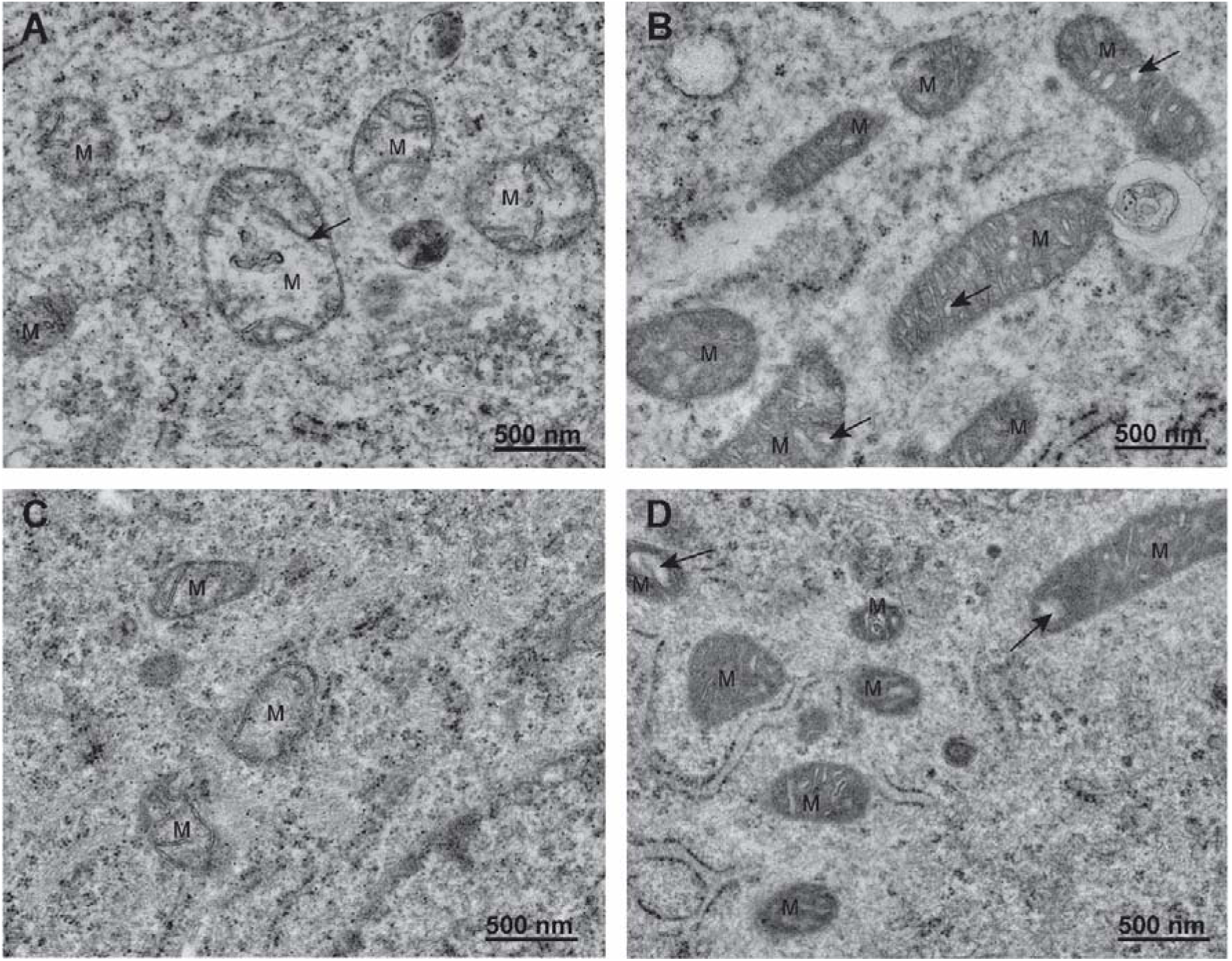
Mitochondria in differentiated GBM cells are smaller and fragmented. **A**) Electron-lucent swollen mitochondria (M) with reduced cristae (arrow) in NIB140 cells. **B**) Condensed electron-dense mitochondria (M) with dilated cristae (arrows) in NIB140 cells. **C**) Electron-lucent swollen mitochondria (M) with reduced cristae in U87 cells. **D**) Condensed electron-dense mitochondria (M) with dilated cristae (arrows) in U87 cells.

### Temozolomide treatment affected the mitochondrial ultrastructure but not the quantity of mitochondria in differentiated GBM cells

After exposure of GBM cells to 48 h treatment with 100 μM TMZ, the ultrastructure of mitochondria was evaluated and quantified. Based on their ultrastructural features, we classified the mitochondria into two following types: normal and defective (Figure **4A**). TMZ treatment induced significant changes in the fractions of mitochondrial types only in differentiated GBM cells (Figures **4A, B**). In both types of differentiated GBM cells there was a significant increase in the fraction of defective mitochondria. In contrast, there were no changes in fraction of defective mitochondria in TMZ-treated GSCs. Despite the significant effect of TMZ treatment on mitochondrial ultrastructure of differentiated GBM cells no effect on the volume density of mitochondria was observed after TMZ treatment in any of cell lines (Supplementary Figure 4C). Structures indicative of autophagy, such as autophagic vacuoles and autophagosomes, were observed both in GSCs and differentiated GBM cells exposed to TMZ (Supplementary Figure **S5**).

**Figure 4.**
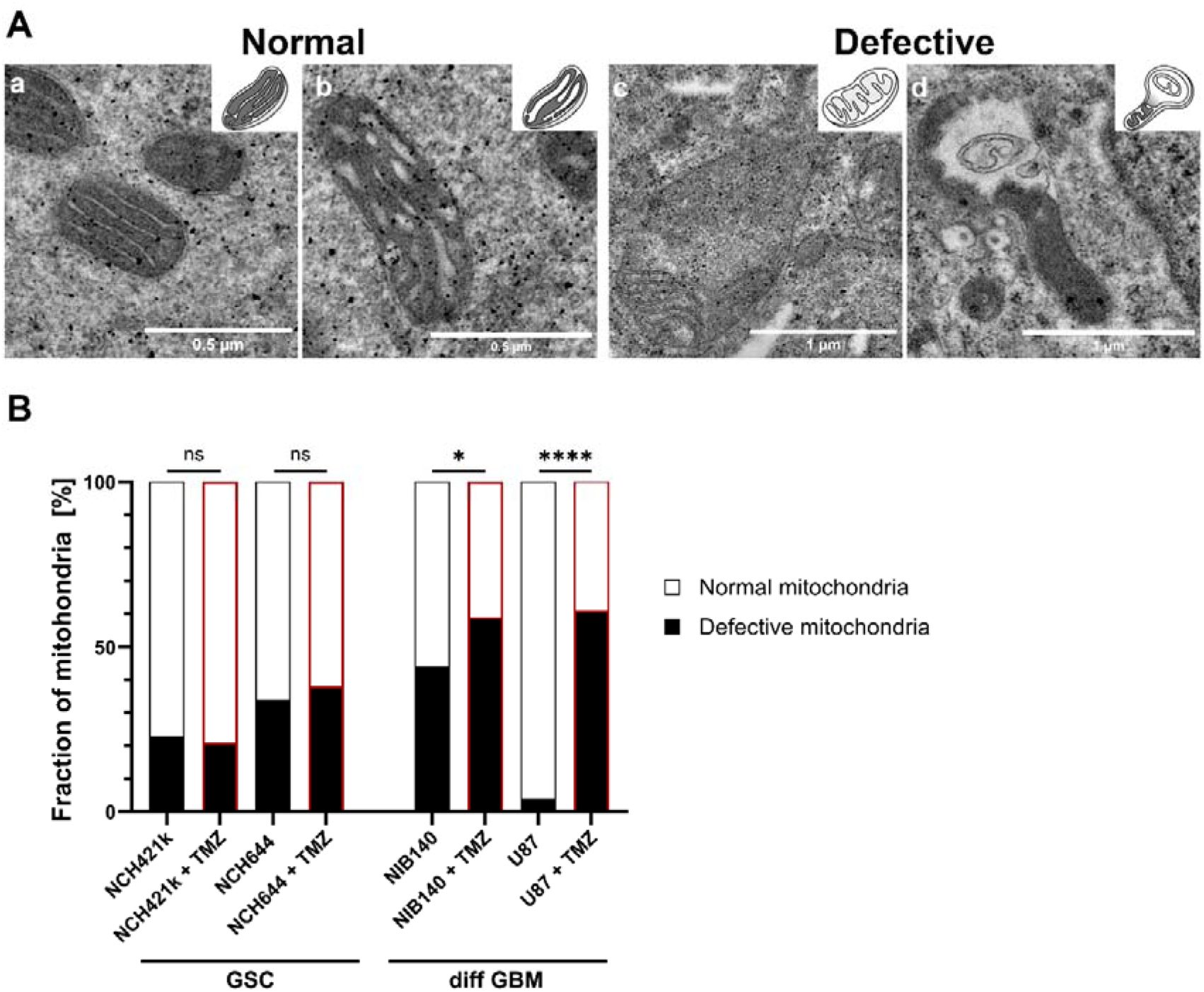
Mitochondrial ultrastructure and its quantification after exposure of cells to 100 µM TMZ. A) Mitochondria were classified into two following types: normal and defective. Normal mitochondria are defined as mitochondria with electron-dense matrix with narrow cristae indicating metabolism with a high OXPHOS rate (a) and mitochondria with electron-dense matrix with dilated cristae indicating metabolism with a lower OXPHOS rate (b). Defective mitochondria are defined as mitochondria with electron-lucent matrix and cristae reduced in size and numbers indicating metabolism without OXPHOS (c) and mitochondria with abnormal ultrastructural alterations (swollen with membrane swirls) indicating damage and stress in mitochondria (d). B) Quantification of the 2 categories of mitochondrial ultrastructure as fraction of all mitochondria. Experiments were performed in two independent repeats (n = 2). Statistics were performed using Fisher’s exact test. * p < 0.05, **** p < 0.0001.

### GSCs are metabolically quiescent and rely more on OXPHOS than differentiated GBM cells

XF analysis was performed to evaluate mitochondrial OXPHOS and glycolysis rates in GSCs NCH421k and NCH644 in comparison to differentiated GBM cells U87 and NIB140 (Figures **5A, B**). Based on ATP production measurements, the metabolic profile of GSCs was quiescent, as they produced less ATP than differentiated GBM cells (Figures **5C, E**). GSCs rely more on OXPHOS than glycolysis for ATP production as the ratio between mitochondrial and glycolytic ATP production rate was higher in GSCs than in differentiated GBM cells (Figures **5D-E**). Glycolysis rates were the highest in differentiated GBM cells U87 and NIB140 and patient-derived cells NIB140 displayed the highest energy production (Figure **5C**).

**Figure 5.**
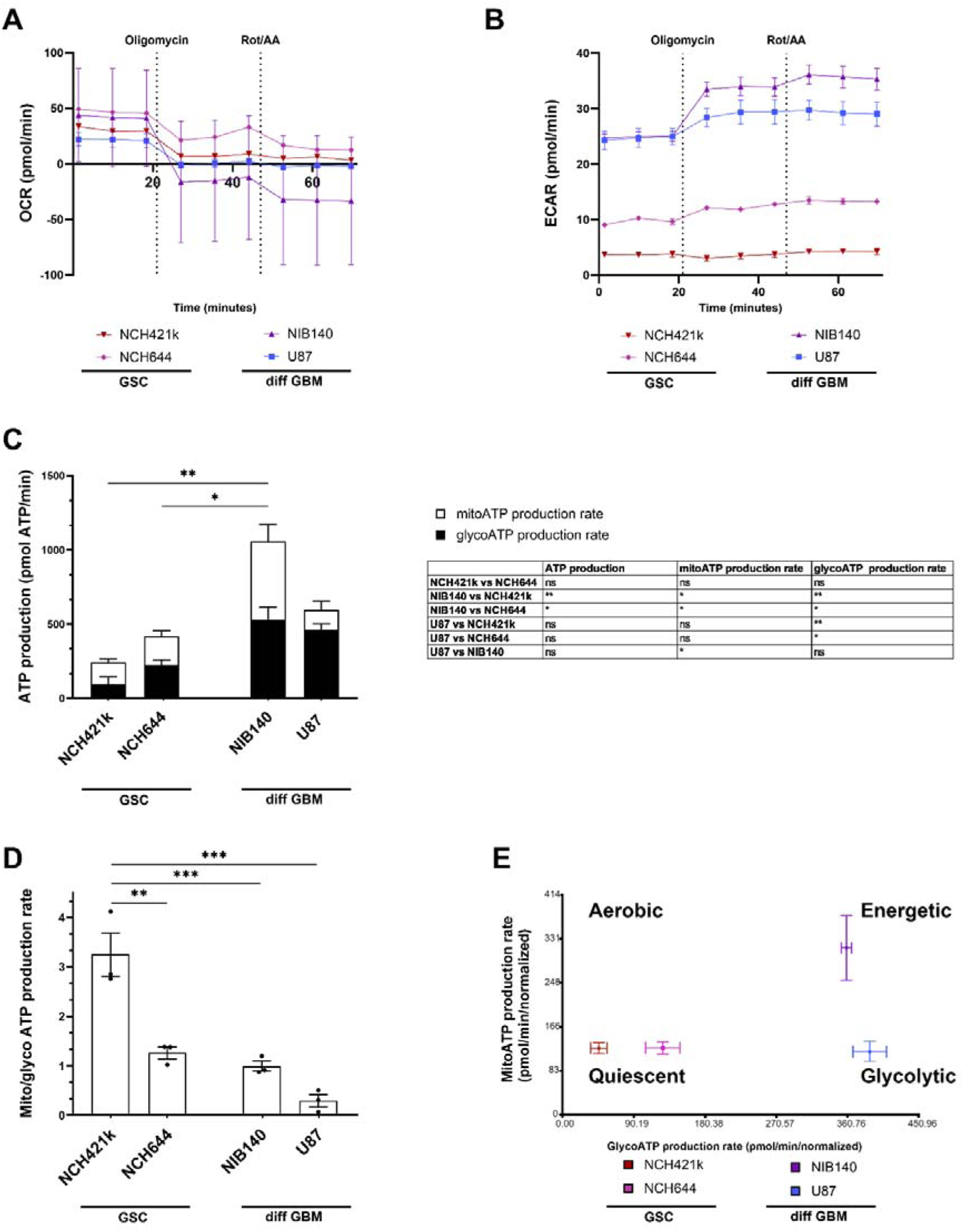
Metabolic profiles of GSCs and differentiated GBM cells as determined by extracellular flux analysis and XF Real-time ATP Rate Assay Kit (Seahorse, Agilent). A) and B) Representative kinetic graphs of oxygen consumption rate (OCR) and extracellular acidification rate (ECAR) in response to oligomycin and rotenone/antimycin A (Rot/AA) are shown. C) Mitochondrial (mito) and glycolytic (glyco) ATP production rate in cells with respective statistics are shown in the table. Statistically significant differences in ATP production are marked directly on the graph. D) Ratio between mito ATP and glyco ATP production rates in cells. E) Metabolic profiles of cells that can be aerobic, energetic, quiescent and glycolytic. Values are shown as means ± SD (A, B) or SEM (C-E). Experiments were performed in 3 independent repeats (n = 3). Statistics were performed using Tukey’s multiple comparisons test. * p < 0.05, ** p < 0.01, *** p < 0.001.

### GSC metabolism relies on OXPHOS after treatment with temozolomide

Cells were treated with 100 µM TMZ for 48 h and after that cellular metabolism was examined. Cell viability and total ATP production were not altered after TMZ treatment, except for the 50% decrease in NCH644 cell viability when treated with the highest concentration of TMZ (Supplementary Figure **S6**). Furthermore, the glycolytic and mitochondrial ATP production was not altered in cells after treatment with TMZ (Figure 6A). We observed that GSCs rely on OXPHOS after TMZ treatment since the ratio between mitochondrial and glycolytic ATP production was above 2 which was much higher than in differentiated GBM cells (Figure 6B). After treatment with TMZ the Glycolytic Rate Assay Kit and Cell Mito Stress Test Kit were performed in cells to evaluate glycolytic rate and mitochondrial function, respectively. Differentiated GBM cells retained high glycolytic ATP production after TMZ treatment. NIB140 cells increased the ECAR rate and compensatory glycolysis after TMZ treatment. There was no change in glycolysis rates in U87, NCH644 and NCH421k cells after TMZ treatment (Supplementary Figure S7). There were also no changes in the monitored parameters of mitochondrial function in cells after treatment with TMZ, except increased maximal respiration in U87 cells after TMZ treatment. NIB140 cells showed the highest proton leak among the cell cultures used (Supplementary Figure S8).

**Figure 6.**
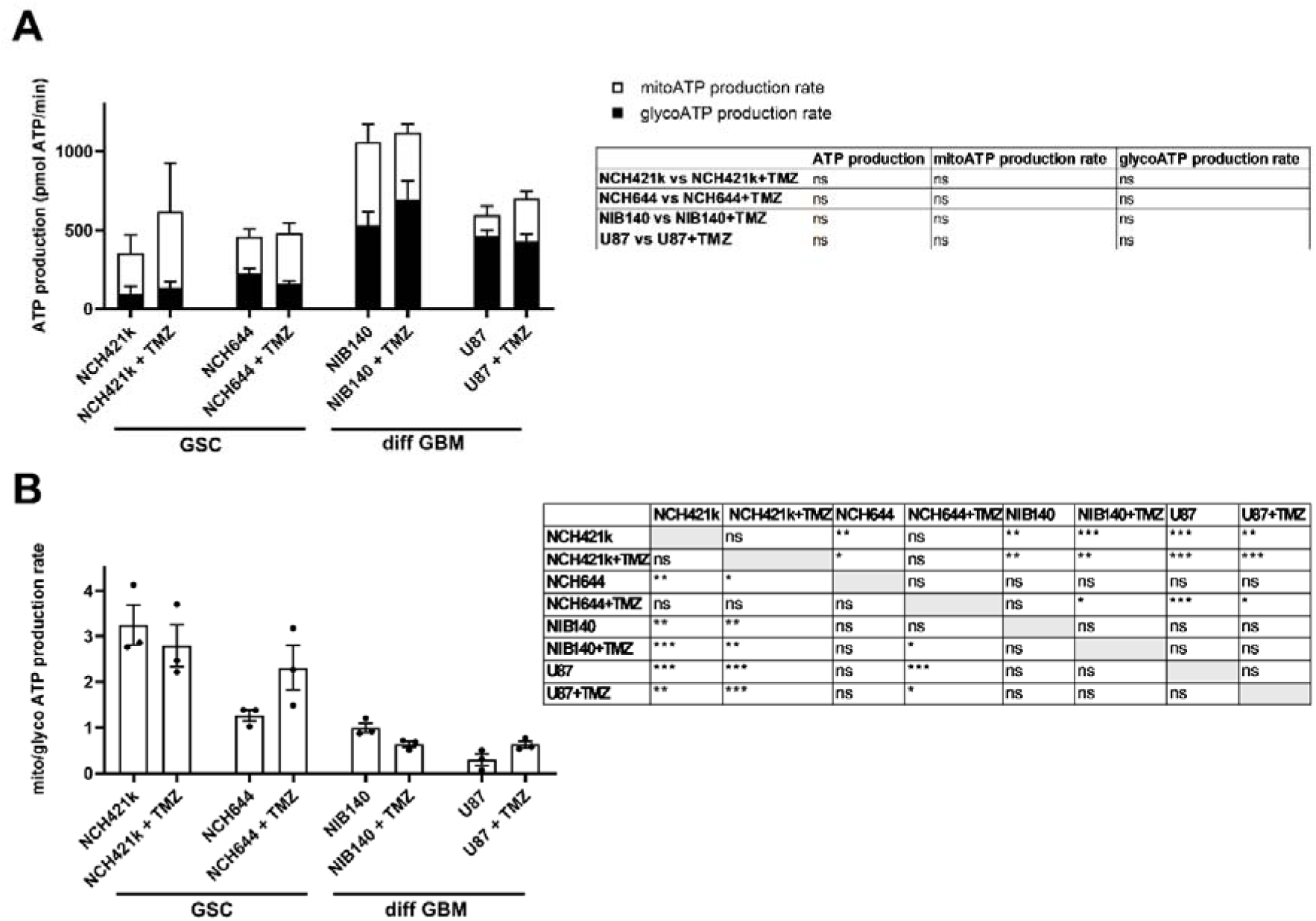
Metabolic profiles and their differences after treatment with 100 µM TMZ in GSCs and differentiated GBM cells and determined by extracellular flux analysis and XF Real-time ATP Rate Assay Kit (Seahorse, Agilent). A) Changes in ATP production, glycolytic (glyco) ATP and mitochondrial (mito) ATP production rate after treatment with TMZ were not detected. B) Differences in ratio between mito and glyco ATP production rates after treatment with TMZ. Values are shown as means ± SEM. Experiments were performed in 3 independent repeats (n = 3). Unpaired t-test was performed to compare statistical difference between control and treated samples within each cell line (A). Tukey’s multiple comparisons test was used to compare all conditions after treatment with TMZ (B). * p < 0.05, ** p < 0.01, *** p < 0.001.

## Discussion

Results of our current study revealed that GSCs have a more quiescent and aerobic metabolic profile than differentiated GBM cells. This is supported both by the microscopic observations of mitochondrial morphology and ultrastructure and by the direct measurements of cellular metabolism by XF analysis.

General ultrastructure showed that GSCs contain little rough endoplasmic reticulum and Golgi apparatus and that their cell nuclei occupy a relatively large fraction of cytoplasm and contain more heterochromatin in comparison to the differentiated GBM cells. These observations are in agreement with the dedifferentiated status of GSCs and their quiescence which is one of important factors in their resistance to therapy [6, 19, 45]. The differentiated GBM cells contain more rough endoplasmic reticulum and Golgi apparatus and nuclei with more euchromatin. This probably supports increased protein synthesis necessary for the excessive growth and proliferation of differentiated GBM cells. The results of the XF analysis support this, showing that GSCs are more quiescent with a lower ATP production rate, while differentiated GBM cells are more energetic, exhibiting a higher ATP production rate. Similar results were reported by Spehalski et al. [46] in their study of GSCs and differentiated GBM cells using XF analysis.

Despite their quiescent phenotype, we observed that the GSCs contained large, elongated mitochondria that occupied a large fraction of the cytoplasm. In contrast, the differentiated GBM cells had small round mitochondria that occupied a smaller portion of the cytoplasm. This indicates enhanced biogenesis and fusion of mitochondria in GSCs and a more fragmented mitochondrial network in differentiated GBM cells. Our XF analysis has shown that the ratio between mitochondrial and glycolytic ATP production is higher in GSCs than in the differentiated GBM cells. Taken together, this shows on cellular and molecular level that the GSCs have a more OXPHOS reliant metabolism, whereas the differentiated GBM cells are more glycolytic. Generally, highly fused mitochondrial networks of elongated mitochondria are associated with high OXPHOS activity particularly in conditions of nutrient withdrawal and mild stress, whereas the fragmented mitochondria indicate impaired OXPHOS and increased glycolysis particularly in conditions of nutrient excess or severe stress that induce apoptosis [24, 47, 48]. These different morphologies of mitochondrial networks in cancer stem cells reflect the differences in tumor microenvironment in which they reside, and the different energetic states related to their differentiation with more quiescent cells favoring fused mitochondria and OXPHOS [16, 26, 49]. In GBM, the heterogeneity of cell types is reflected in the heterogeneity of their metabolic profiles [50]. The results of our current study are in line with previous reports [5, 20].

Observed differences in metabolic profiles between GSCs and differentiated GBM cells were also evident in the ultrastructure of mitochondria, particularly in the morphology of cristae. GSCs contained structurally intact mitochondria. Most of GSCs contained electron-dense mitochondria with narrow cristae corresponding with high OXPHOS activity. In a smaller subset of GSCs, the mitochondria were electron-dense with dilated cristae corresponding with reduced OXPHOS activity. In contrast, the differentiated GBM cells contained mitochondria with varying ultrastructural features, with the two most common types being electron-dense mitochondria with dilated cristae, which corresponded with reduced OXPHOS activity and the electron-lucent swollen mitochondria with reduced cristae, which associated with impaired OXPHOS activity. Previous reports on brain tumor cell ultrastructure, including GBM cells, also describe electron-dense mitochondria with dilated cristae and electron-lucent swollen mitochondria [30, 31, 53]. The electron-lucent swollen mitochondria with reduced cristae generally associate with the mitochondrial damage due to Ca^2+^ overload and excessive ROS production [62–64] and have impaired OXPHOS activity [15, 58].

Another interesting observation in our investigation is the presence of cristae that are oriented along the longitudinal axis of the mitochondria. This was particularly prominent in GSCs, where nearly all mitochondria exhibited this orientation of cristae. The important regulators of cristae shape are the mitochondrial contact sites and cristae-organizing system (MICOS) complex, ATP synthase complex, optic atrophy 1 (OPA1) protein and lipid composition of the inner mitochondrial membrane [48, 57]. In yeast, mutations in genes that encode certain subunits of ATP synthase and components of the MICOS complex result in mitochondrial cristae oriented along the long axis of mitochondria [65–67]. Thus, reduced or even absent amounts of longitudinal oriented cristae in differentiated GBM cell mitochondria may well indicate that the GBM cells exhibit defects in their ATP synthase or MICOS complex.

The observed high mitochondrial content, intact ultrastructural integrity of mitochondria and high ratio between mitochondrial and glycolytic ATP production in GSCs are in agreement with the findings of previous studies, which employed various metabolic assays and have shown that the GSC are less glycolytic than the differentiated GBM cells [5, 17, 19]. The GSCs are probably capable of surviving in hypoxic microenvironment due to their relatively quiescent state [6, 16, 19], which requires a relatively low production of ATP. Dilated cristae observed in mitochondria of differentiated GBM cells are associated with increased ROS production due to the less effective electron transport chain [68]. The increased production of ROS is known to induce certain signaling pathways that increase proliferation and survival of cancer cells [14, 17, 21, 54] and are thus crucial in tumorigenesis. The relatively low mitochondrial content, the fragmented shape of the mitochondrial network and the presence of swollen mitochondria indicate that the differentiated GBM are heavily dependent on glycolysis. It is known that numerous cancer cells rely on glycolysis even in the presence of oxygen due to defective mitochondria [13–15]. The glycolysis provides highly proliferative cancer cells with high ATP production under low oxygen conditions. Besides this, it supplies the precursors for anabolic pathways and abundant NADPH for reductive biosynthetic reactions and antioxidant defenses [6, 21, 22, 69].

Our results indicate that treatment with chemotherapeutic TMZ that is used in standard treatment procedures for GBM patients induces significant ultrastructural mitochondrial damage in the differentiated GBM cells but has less effect on the ultrastructure of mitochondria in GSCs. However, the observed ultrastructural mitochondrial damage in the differentiated GBM cells after TMZ treatment is not reflected in the metabolic profile or cell viability. TMZ is a DNA alkylating agent. The DNA damage caused by the TMZ has been shown to induce increased ROS production in mitochondria which contributes to the apoptosis of susceptible GBM cells [70, 71]. Apoptosis is in a large part mediated by mitochondria, including the increased ROS generation in mitochondria, increased mitochondrial membrane permeability and finally the structural damage of mitochondria [72]. The mitochondria thus have a critical role in GBM resistance to TMZ treatment [73]. It has been shown that the TMZ-resistant GBM cells reorganize the electron transport chain in their mitochondria towards more efficient mitochondrial coupling and decreased ROS production [74, 75]. This may well explain the decreased ultrastructural mitochondrial damage in GSCs in comparison to the differentiated GBM cells observed in our study after TMZ treatment.

Taken together, we showed differences in mitochondria ultrastructure and cellular metabolism between stem-like GSCs and differentiated GBM cells in normal conditions and upon chemotherapy. TMZ treatment induced significant ultrastructural mitochondrial damage in differentiated GBM cells and had less effect on mitochondria in GSCs, indicating that the mitochondria play an important role in GSC resistance to TMZ treatment. Although further studies are required to fully understand GBM therapeutic resistance and the effects of different treatment regimens on the mitochondrial structure and function of GSCs, this study establishes a foundation for a deeper understanding of the metabolic heterogeneity of glioblastoma cells, including both stem and differentiated cells, and their roles in therapy response and resistance.

## Funding and Acknowledgements

We would like to thank Dhaval Patel from the Health Science Center at the Texas Tech University, Amarillo, USA, for technical assistance with Seahorse experiments. This work was supported by the Slovenian Research and Innovation Agency (ARIS) grants (P1-0245 (B.B., M.N.), J3-4504 (M.N.), BI-US/22-24-007 (B.B), J3-2526 (C.J.F.V.N.), BI-US/24-26-022 (M.N.), Young Researcher grant (SKM; 59639), by University Infrastructural Centre “Microscopy of Biological Samples” at the Biotechnical Faculty, University of Ljubljana (M.V.), Texas Tech University start-up (to K.F.T.), Cancer Prevention and Research Institute of Texas Scholar Award RR200059 (to K.F.T.), the Foundation for Prader–Willi Syndrome Research Grant 22-0321 and 23-0447 (to K.F.T.), Fulbright Fellowship Scholar (B.B.) and European Union’s Horizon project Twinning for excellence to strategically advance research in carcinogenesis and cancer (CutCancer; 101079113) (B.B. and M.N.).

## Authorship

UB, MN, MV, CJFVN and BB designed the study. UB, MN, SKG, KFT, MV, CJFVN and BB designed and performed experiments and analysed and interpreted data. UB, MV, CJFVN, and BB wrote the first draft of the manuscript. All authors reviewed and approved the submitted manuscript.

## Data Availability

All data generated or analysed during this study are included in this published paper and its Supplementary File. The datasets generated and analysed during the current study are also available from the corresponding author upon reasonable request.

## References

1. van Solinge TS, Nieland L, Chiocca EA, Broekman MLD. Advances in local therapy for glioblastoma - taking the fight to the tumour. Nat Rev Neurol. 2022;18:221–36.

2. Stupp R, Hegi ME, Mason WP, van den Bent MJ, Taphoorn MJ, Janzer RC, et al. Effects of radiotherapy with concomitant and adjuvant temozolomide versus radiotherapy alone on survival in glioblastoma in a randomised phase III study: 5-year analysis of the EORTC-NCIC trial. Lancet Oncol. 2009;10:459–66.

3. Neftel C, Laffy J, Filbin MG, Hara T, Shore ME, Rahme GJ, et al. An Integrative Model of Cellular States, Plasticity, and Genetics for Glioblastoma. Cell. 2019;178:835–849.e21.

4. Prager BC, Bhargava S, Mahadev V, Hubert CG, Rich JN. Glioblastoma Stem Cells: Driving Resilience through Chaos. Trends in Cancer. 2020;6:223–35.

5. Vlashi E, Lagadec C, Vergnes L, Matsutani T, Masui K, Poulou M, et al. Metabolic state of glioma stem cells and nontumorigenic cells. Proc Natl Acad Sci U S A. 2011;108:16062–7.

6. Badr CE, Silver DJ, Siebzehnrubl FA, Deleyrolle LP. Metabolic heterogeneity and adaptability in brain tumors. Cell Mol Life Sci. 2020;77:5101–19.

7. Hira VVV, Breznik B, Vittori M, Loncq de Jong A, Mlakar J, Oostra RJ, et al. Similarities Between Stem Cell Niches in Glioblastoma and Bone Marrow: Rays of Hope for Novel Treatment Strategies. J Histochem Cytochem. 2020;68:33–57.

8. Hira VVV, Aderetti DA, van Noorden CJF. Glioma Stem Cell Niches in Human Glioblastoma Are Periarteriolar. Journal of Histochemistry and Cytochemistry. 2018;66:349–58.

9. Schiffer D, Annovazzi L, Casalone C, Corona C, Mellai M. Glioblastoma: Microenvironment and niche concept. Cancers. 2019;11.

10. Li P, Zhou C, Xu L, Xiao H. Hypoxia enhances stemness of cancer stem cells in glioblastoma: an in vitro study. Int J Med Sci. 2013;10:399–407.

11. Heddleston JM, Li Z, McLendon RE, Hjelmeland AB, Rich JN. The hypoxic microenvironment maintains glioblastoma stem cells and promotes reprogramming towards a cancer stem cell phenotype. Cell Cycle. 2009;8:3274–84.

12. Iranmanesh Y, Jiang B, Favour OC, Dou Z, Wu J, Li J, et al. Mitochondria’s Role in the Maintenance of Cancer Stem Cells in Glioblastoma. Front Oncol. 2021;11.

13. Carew JS, Huang P. Mitochondrial defects in cancer. Mol Cancer. 2002;1.

14. Wallace DC. Mitochondria and cancer: Warburg addressed. Cold Spring Harb Symp Quant Biol. 2005;70:363–74.

15. Seyfried TN, Flores RE, Poff AM, D’Agostino DP. Cancer as a metabolic disease: implications for novel therapeutics. Carcinogenesis. 2014;35:515–27.

16. Peiris-Pagès M, Martinez-Outschoorn UE, Pestell RG, Sotgia F, Lisanti MP. Cancer stem cell metabolism. Breast Cancer Res. 2016;18.

17. Strickland M, Stoll EA. Metabolic Reprogramming in Glioma. Front cell Dev Biol. 2017;5 APR.

18. van Noorden CJF, Yetkin-Arik B, Serrano Martinez P, Bakker N, van Breest Smallenburg ME, Schlingemann RO, et al. New Insights in ATP Synthesis as Therapeutic Target in Cancer and Angiogenic Ocular Diseases. J Histochem Cytochem. 2024;72:329–52.

19. van Noorden CJF, Breznik B, Novak M, van Dijck AJ, Tanan S, Vittori M, et al. Cell Biology Meets Cell Metabolism: Energy Production Is Similar in Stem Cells and in Cancer Stem Cells in Brain and Bone Marrow. J Histochem Cytochem. 2022;70:29–51.

20. van Noorden CJF, Hira VVV, van Dijck AJ, Novak M, Breznik B, Molenaar RJ. Energy Metabolism in IDH1 Wild-Type and IDH1-Mutated Glioblastoma Stem Cells: A Novel Target for Therapy? Cells. 2021;10:1–16.

21. Wallace DC. Mitochondria and cancer. Nat Rev Cancer. 2012;12:685–98.

22. Vyas S, Zaganjor E, Haigis MC. Mitochondria and Cancer. Cell. 2016;166:555–66.

23. Ordys BB, Launay S, Deighton RF, McCulloch J, Whittle IR. The role of mitochondria in glioma pathophysiology. Mol Neurobiol. 2010;42:64–75.

24. Wai T, Langer T. Mitochondrial Dynamics and Metabolic Regulation. Trends Endocrinol Metab. 2016;27:105–17.

25. Senft D, Ronai ZA. Regulators of mitochondrial dynamics in cancer. Curr Opin Cell Biol. 2016;39:43–52.

26. Trotta AP, Chipuk JE. Mitochondrial dynamics as regulators of cancer biology. Cell Mol Life Sci. 2017;74:1999–2017.

27. Maycotte P, Marín-Hernández A, Goyri-Aguirre M, Anaya-Ruiz M, Reyes-Leyva J, Cortés-Hernández P. Mitochondrial dynamics and cancer. Tumour Biol. 2017;39.

28. García-Heredia JM, Carnero A. Role of Mitochondria in Cancer Stem Cell Resistance. Cells. 2020;9.

29. De Luca A, Fiorillo M, Peiris-Pagès M, Ozsvari B, Smith DL, Sanchez-Alvarez R, et al. Mitochondrial biogenesis is required for the anchorage-independent survival and propagation of stem-like cancer cells. Oncotarget. 2015;6:14777–95.

30. Arismendi-Morillo GJ, Castellano-Ramirez A V. Ultrastructural mitochondrial pathology in human astrocytic tumors: potentials implications pro-therapeutics strategies. J Electron Microsc (Tokyo). 2008;57:33–9.

31. Arismendi-Morillo G. Electron microscopy morphology of the mitochondrial network in gliomas and their vascular microenvironment. Biochim Biophys Acta. 2011;1807:602–8.

32. Scheithauer BW, Bruner JM. The ultrastructural spectrum of astrocytic neoplasms. Ultrastruct Pathol. 1987;11:535–81.

33. Liberski PP, Kordek R. Ultrastructural pathology of glial brain tumors revisited: a review. Ultrastruct Pathol. 1997;21:1–31.

34. Sipe, JC, Herman, MM RL. Electron microscopic observations on human glioblastomas and astrocytomas maintained in organ culture systems. Am J Pathol. 1973;73:589–606.

35. Khurshed M, Aarnoudse N, Hulsbos R, Hira VVV, Van Laarhoven HWM, Wilmink JW, et al. IDH1-mutant cancer cells are sensitive to cisplatin and an IDH1-mutant inhibitor counteracts this sensitivity. FASEB J. 2018;32:6344–52.

36. Verberk SGS, de Goede KE, Gorki FS, van Dierendonck XAMH, Argüello RJ, Van den Bossche J. An integrated toolbox to profile macrophage immunometabolism. Cell reports methods. 2022;2.

37. Breznik B, Ko MW, Tse C, Chen PC, Senjor E, Majc B, et al. Infiltrating natural killer cells bind, lyse and increase chemotherapy efficacy in glioblastoma stem-like tumorospheres. Commun Biol. 2022;5.

38. Porčnik A, Novak M, Breznik B, Majc B, Hrastar B, Šamec N, et al. TRIM28 Selective Nanobody Reduces Glioblastoma Stem Cell Invasion. Molecules. 2021;26.

39. Majc B, Habič A, Novak M, Rotter A, Porčnik A, Mlakar J, et al. Upregulation of Cathepsin X in Glioblastoma: Interplay with γ-Enolase and the Effects of Selective Cathepsin X Inhibitors. Int J Mol Sci. 2022;23.

40. Novak M, Majc B, Malavolta M, Porčnik A, Mlakar J, Hren M, et al. The Slovenian translational platform GlioBank for brain tumour research: identification of molecular signatures of glioblastoma progression. Neuro-Oncology Adv. 2025. 10.1093/NOAJNL/VDAF015.

41. Majc B, Habič A, Malavolta M, Vittori M, Porčnik A, Bošnjak R, et al. Patient-derived tumor organoids mimic treatment-induced DNA damage response in glioblastoma. iScience. 2024;27.

42. Breznik B, Motaln H, Vittori M, Rotter A, Turnšek TL. Mesenchymal stem cells differentially affect the invasion of distinct glioblastoma cell lines. Oncotarget. 2017;8:25482–99.

43. Schindelin J, Arganda-Carreras I, Frise E, Kaynig V, Longair M, Pietzsch T, et al. Fiji: an open-source platform for biological-image analysis. Nat Methods. 2012;9:676–82.

44. Cardona A, Saalfeld S, Schindelin J, Arganda-Carreras I, Preibisch S, Longair M, et al. TrakEM2 software for neural circuit reconstruction. PLoS One. 2012;7.

45. Lah TT, Novak M, Breznik B. Brain malignancies: Glioblastoma and brain metastases. Semin Cancer Biol. 2020;60:262–73.

46. Spehalski EI, Lee JA, Peters C, Tofilon P, Camphausen K. The Quiescent Metabolic Phenotype of Glioma Stem Cells. J Proteomics Bioinform. 2019;12.

47. Galloway CA, Lee H, Yoon Y. Mitochondrial morphology-emerging role in bioenergetics. Free Radic Biol Med. 2012;53:2218–28.

48. Glancy B, Kim Y, Katti P, Willingham TB. The Functional Impact of Mitochondrial Structure Across Subcellular Scales. Front Physiol. 2020;11.

49. Peixoto J, Lima J. Metabolic traits of cancer stem cells. Dis Model Mech. 2018;11.

50. Duraj T, García-romero N, Carrión-navarro J, Madurga R, de Mendivil AO, Prat-acin R, et al. Beyond the Warburg Effect: Oxidative and Glycolytic Phenotypes Coexist within the Metabolic Heterogeneity of Glioblastoma. Cells. 2021;10:1–23.

51. Mao P, Joshi K, Li J, Kim SH, Li P, Santana-Santos L, et al. Mesenchymal glioma stem cells are maintained by activated glycolytic metabolism involving aldehyde dehydrogenase 1A3. Proc Natl Acad Sci U S A. 2013;110:8644–9.

52. Shibao S, Minami N, Koike N, Fukui N, Yoshida K, Saya H, et al. Metabolic heterogeneity and plasticity of glioma stem cells in a mouse glioblastoma model. Neuro Oncol. 2018;20:343–54.

53. Arismendi-Morillo G, Castellano-Ramírez A, Seyfried TN. Ultrastructural characterization of the Mitochondria-associated membranes abnormalities in human astrocytomas: Functional and therapeutics implications. Ultrastruct Pathol. 2017;41:234–44.

54. Zick M, Rabl R, Reichert AS. Cristae formation-linking ultrastructure and function of mitochondria. Biochim Biophys Acta. 2009;1793:5–19.

55. Hackenbrock CR. Ultrastructural bases for metabolically linked mechanical activity in mitochondria. II. Electron transport-linked ultrastructural transformations in mitochondria. J Cell Biol. 1968;37:345–69.

56. Hackenbrock CR. Ultrastructural bases for metabolically linked mechanical activity in mitochondria. I. Reversible ultrastructural changes with change in metabolic steady state in isolated liver mitochondria. J Cell Biol. 1966;30:269–97.

57. Kondadi AK, Anand R, Reichert AS. Cristae Membrane Dynamics - A Paradigm Change. Trends Cell Biol. 2020;30:923–36.

58. Moscheni C, Malucelli E, Castiglioni S, Procopio A, De Palma C, Sorrentino A, et al. 3D Quantitative and Ultrastructural Analysis of Mitochondria in a Model of Doxorubicin Sensitive and Resistant Human Colon Carcinoma Cells. Cancers (Basel). 2019;11.

59. Ježek P, Jabůrek M, Holendová B, Engstová H, Dlasková A. Mitochondrial Cristae Morphology Reflecting Metabolism, Superoxide Formation, Redox Homeostasis, and Pathology. Antioxid Redox Signal. 2023;39:635–83.

60. Dlasková A, Špaček T, Engstová H, Špačková J, Schröfel A, Holendová B, et al. Mitochondrial cristae narrowing upon higher 2-oxoglutarate load. Biochim Biophys acta Bioenerg. 2019;1860:659–78.

61. Alirol E, Martinou JC. Mitochondria and cancer: is there a morphological connection? Oncogene. 2006;25:4706–16.

62. Galindo MF, Jordán J, González-García C, Ceña V. Reactive oxygen species induce swelling and cytochrome c release but not transmembrane depolarization in isolated rat brain mitochondria. Br J Pharmacol. 2003;139:797–804.

63. Peng T-I, Jou M-J. Mitochondrial swelling and generation of reactive oxygen species induced by photoirradiation are heterogeneously distributed. Ann N Y Acad Sci. 2004;1011:112–22.

64. Kaasik A, Safiulina D, Zharkovsky A, Veksler V. Regulation of mitochondrial matrix volume. Am J Physiol Cell Physiol. 2007;292.

65. P P, J V, B C, J S, V S, DM M, et al. The ATP synthase is involved in generating mitochondrial cristae morphology. EMBO J. 2002;21:221–30.

66. Stephan T, Brüser C, Deckers M, Steyer AM, Balzarotti F, Barbot M, et al. MICOS assembly controls mitochondrial inner membrane remodeling and crista junction redistribution to mediate cristae formation. EMBO J. 2020;39.

67. Klecker T, Westermann B. Pathways shaping the mitochondrial inner membrane. Open Biol. 2021;11.

68. Plecitá-Hlavatá L, Ježek P. Integration of superoxide formation and cristae morphology for mitochondrial redox signaling. Int J Biochem Cell Biol. 2016;80:31–50.

69. Vaupel P, Multhoff G. Revisiting the Warburg effect: historical dogma versus current understanding. J Physiol. 2021;599:1745–57.

70. Campos-Sandoval JA, Gómez-García MC, Santos-Jiménez J de los, Matés JM, Alonso FJ, Márquez J. Antioxidant responses related to temozolomide resistance in glioblastoma. Neurochem Int. 2021;149.

71. Zhang W Bin, Wang Z, Shu F, Jin YH, Liu HY, Wang QJ, et al. Activation of AMP-activated protein kinase by temozolomide contributes to apoptosis in glioblastoma cells via p53 activation and mTORC1 inhibition. J Biol Chem. 2010;285:40461–71.

72. Fleury C, Mignotte B, Vayssière JL. Mitochondrial reactive oxygen species in cell death signaling. Biochimie. 2002;84:131–41.

73. Li HY, Feng YH, Lin CL, Hsu TI. Mitochondrial Mechanisms in Temozolomide Resistance: Unraveling the Complex Interplay and Therapeutic Strategies in Glioblastoma. Mitochondrion. 2024;75.

74. Oliva CR, Nozell SE, Diers A, McClugage SG, Sarkaria JN, Markert JM, et al. Acquisition of temozolomide chemoresistance in gliomas leads to remodeling of mitochondrial electron transport chain. J Biol Chem. 2010;285:39759–67.

75. Oliva CR, Moellering DR, Gillespie GY, Griguer CE. Acquisition of chemoresistance in gliomas is associated with increased mitochondrial coupling and decreased ROS production. PLoS One. 2011;6.

