## Supplementary File for "Heterogenous mitochondrial ultrastructure and metabolism of human glioblastoma cells: differences between stem-like and differentiated cancer cells in response to chemotherapy"

By authors

Urban Bogataj, Metka Novak, Katrin Galun, Klementina Fon Tacer, Miloš Vittori, Cornelis J.F. Van Noorden, Barbara Breznik

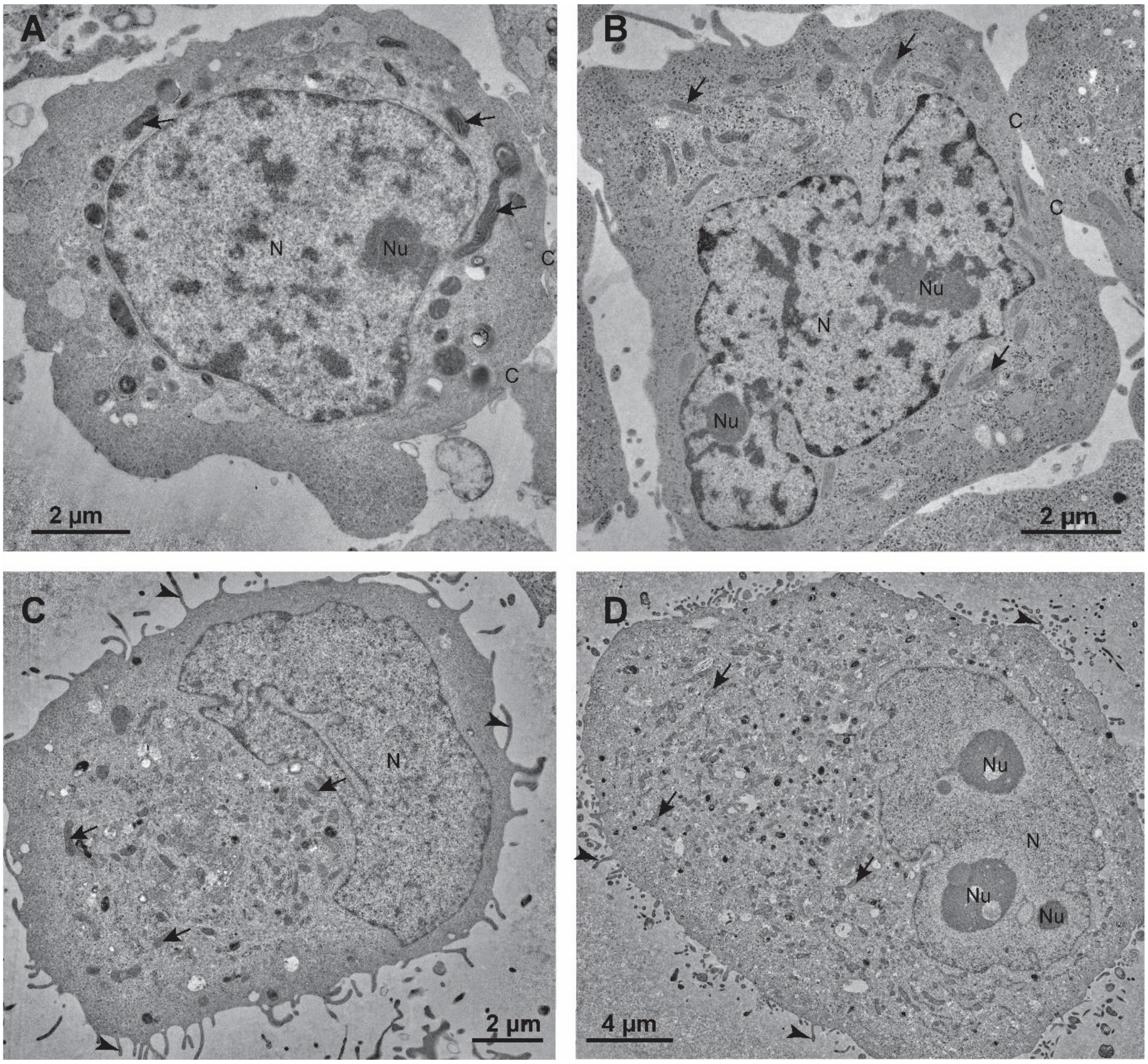


**Supplementary Figure S1:** General ultrastructure of GSCs and differentiated GBM cells. **A**) GSCs NCH421k and **B**) GSCs NCH644 have cell nuclei (N) with abundant heterochromatin. GSCs are in contact with neighboring cells through distinct contact (C) sites. **C**) Differentiated GBM cells NIB140 and **D**) differentiated GBM cells U87 have cell nuclei (N) almost without heterochromatin and exhibit microvilli like projections (arrowheads) at their surface. Arrows – mitochondria, Nu – nucleoli.

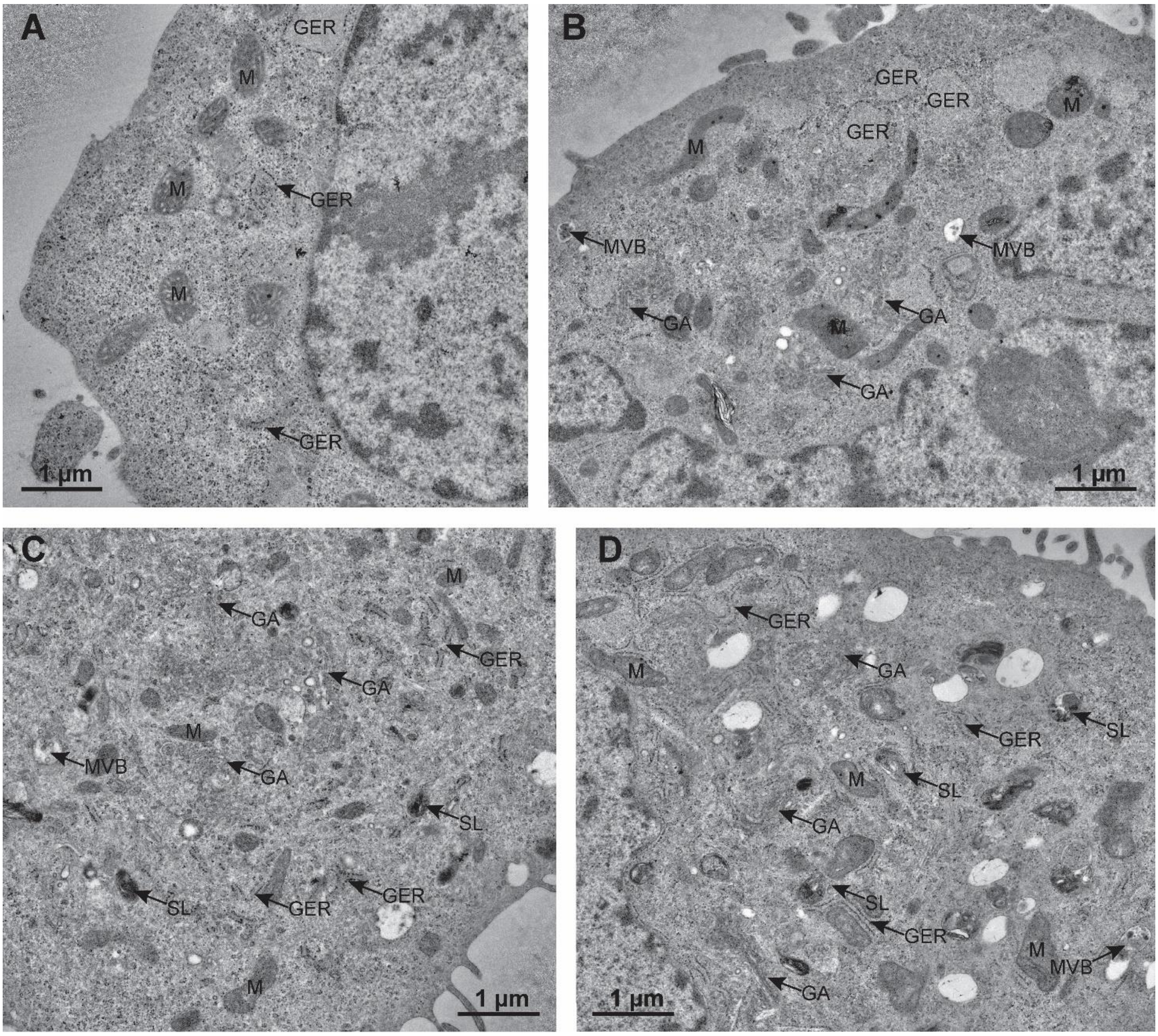


**Supplementary Figure S2:** Endoplasmic reticulum and Golgi apparatus in GSCs and differentiated GBM cells. **A)** GSCs NCH421k and **B)** GSCs NCH644 contain scarce cisternae of rough endoplasmic reticulum (GER) and occasional stacks of Golgi apparatus (GA). **C)** Differentiated GBM cells NIB 140 and **D)** differentiated GBM cells U87 contain abundant cisternae of rough endoplasmic reticulum (GER) and numerous Golgi stacks (GA). M – mitochondria, MVB – multivesicular bodies, SL – secondary lysosomes.

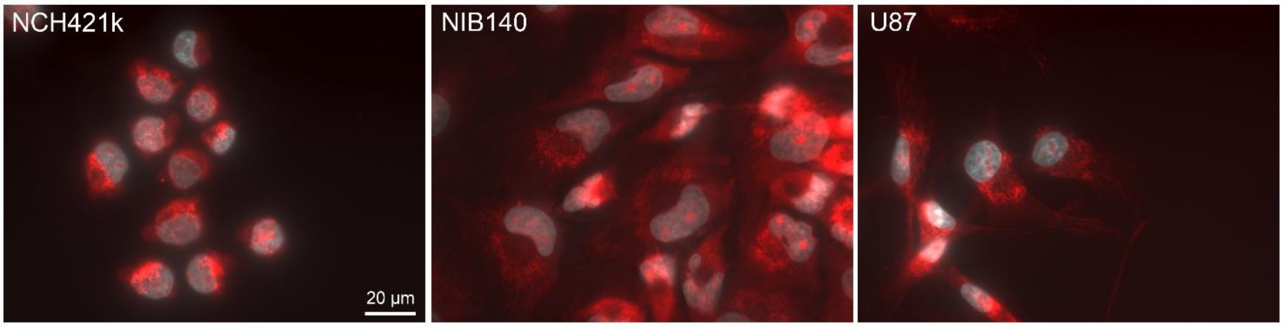


**Supplementary Figure S3:** MitoTracker staining of GSCs and differentiated GBM cells showing the density and distribution of active mitochondria in the cells.

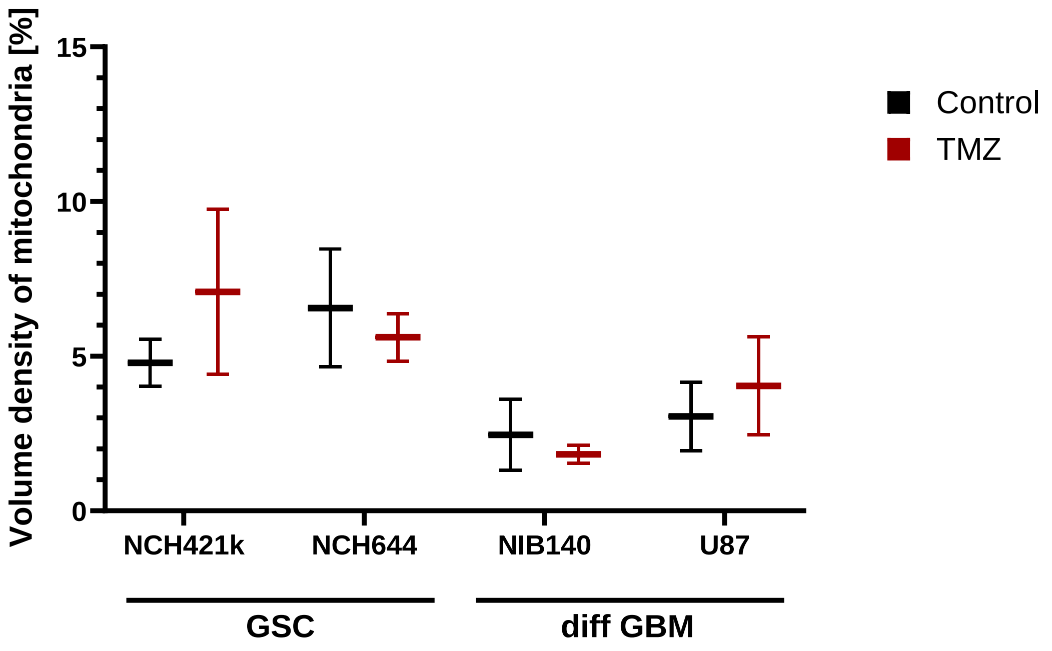


**Supplementary Figure S4:** Volume density of mitochondria showing no effects of treatment with 100 µM TMZ. Experiments were performed in two independent repeats (n = 2). Statistics were performed using an unpaired t-test.

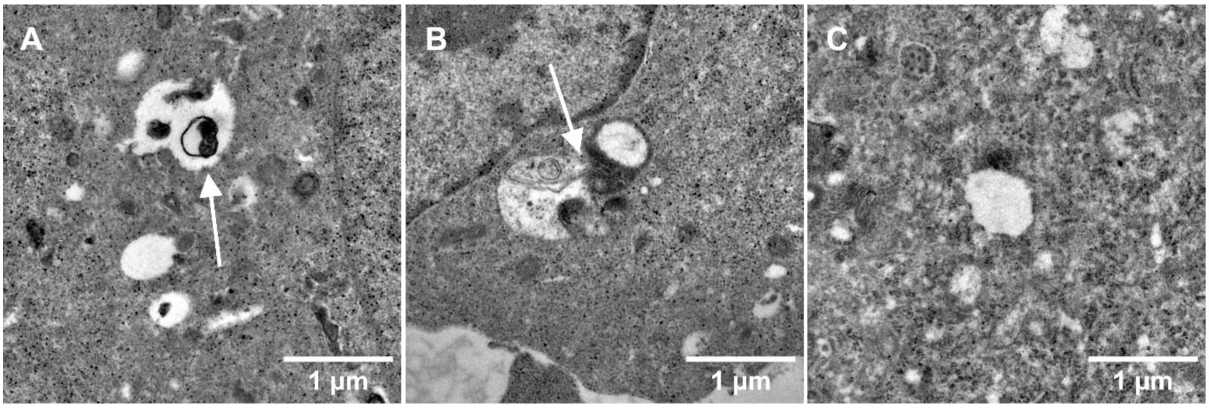


**Supplementary Figure S5:** Examples of autophagy in differentiated GBM cells NIB140 and GSCs NCH644 after TMZ treatment. Images show structures resembling **A**) autophagic vacuole, **B**) contact between swollen mitochondrion and autophagic vacuole, and **C**) autophagosome. Arrow – autophagic vacuole-like structures.

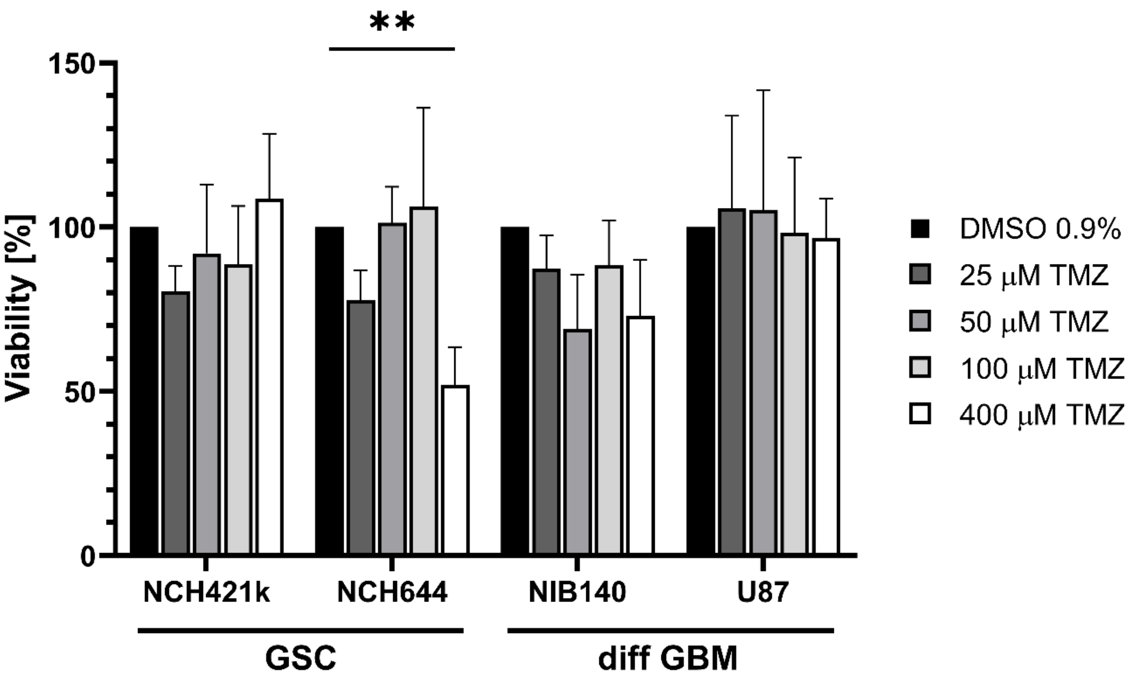


**Supplementary Figure S6:** Viability of GSCs and differentiated GBM cells after TMZ treatment. The cells were treated with a sseries of TMZ concentrations and cell viability was evaluated after 48 h TMZ treatment using MTT and MTS assays. MTT was performed for attached differentiated GBM cells and MTS for floating GSCs. The effect of TMZ was compared with vehicle control (0.9% DMSO). Experiments were performed in three independent repeats (n = 3). One-way ANOVA was performed to compare statistical differences between vehicle control and TMZ-treated samples within each cell line. ** $p < 0.01$.

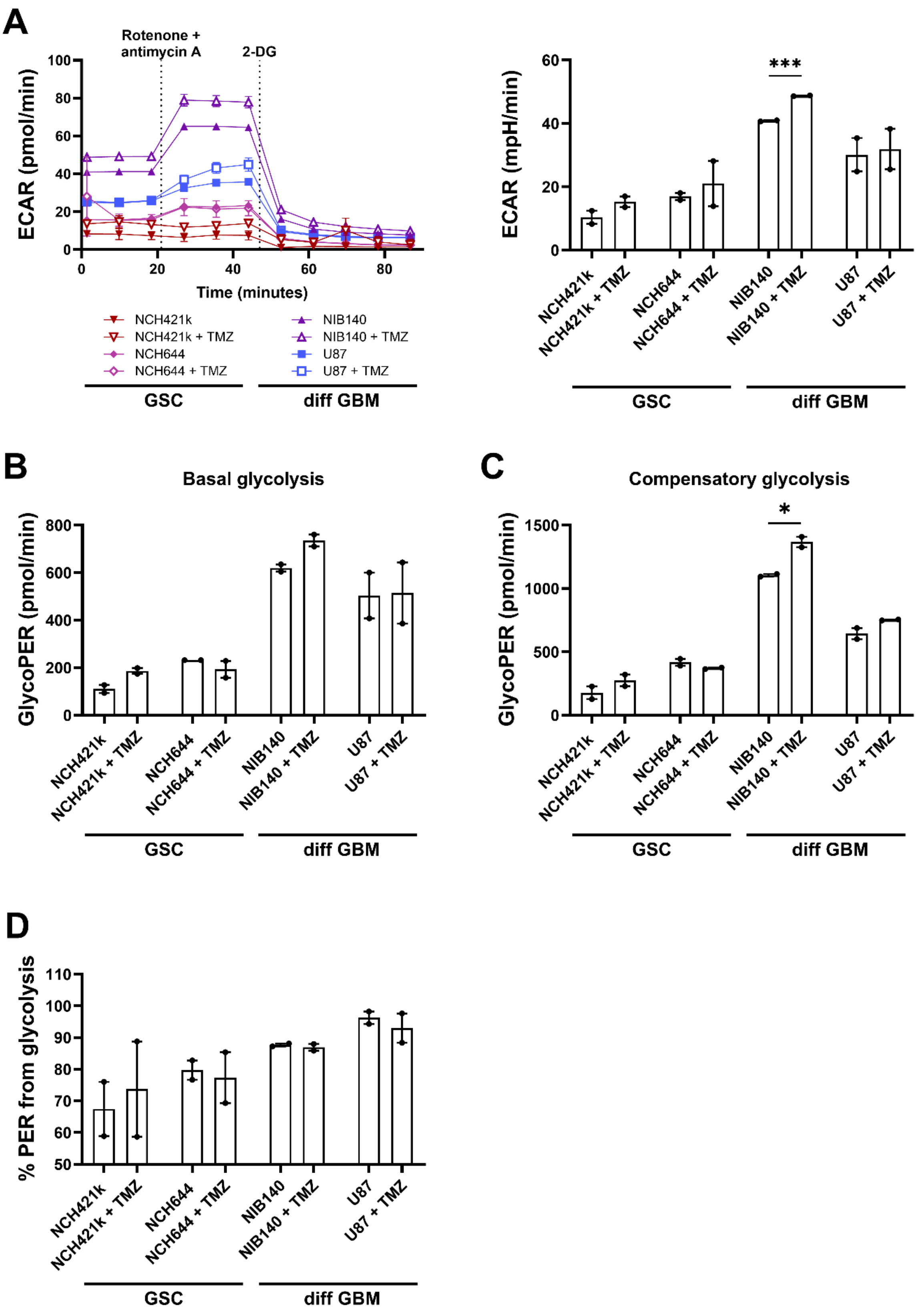


**Supplementary Figure S7.** Glycolysis rates in GSCs and differentiated GBM cells after treatment with TMZ as determined by extracellular flux analysis and XF Glycolytic Rate Assay Kit (Seahorse, Agilent).

A) Kinetic graph of extracellular acidification rates (ECAR), B) basal glycolysis expressed as glycolytic proton efflux rate derived from glycolysis (glycoPER), C) compensatory glycolysis expressed as glycoPER and D) the contribution of glycolysis to PER (% of PER from glycolysis). Values are shown as means ± SEM. Experiments were performed in two independent repeats ($n = 2$). Unpaired t-test was performed to compare statistical differences between control and treated samples within each cell line (A-C). Tukey's multiple comparisons test was used to compare all conditions after treatment with TMZ (D). * $p < 0.05$, *** $p < 0.001$. Abbreviations: 2-DG, 2-deoxyglucose; PER, Proton efflux rate.

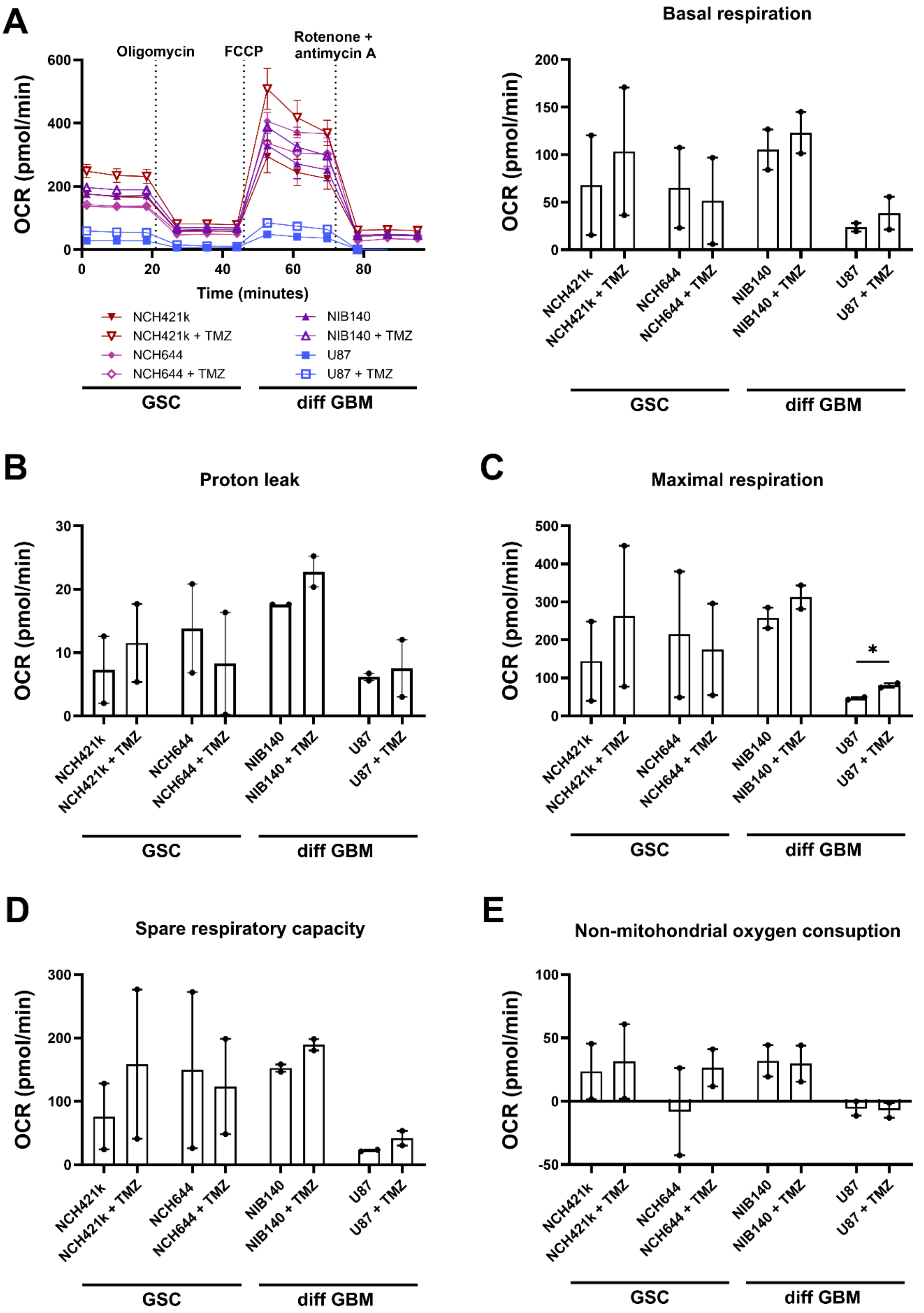


**Supplementary Figure S8.** Mitochondrial respiration rates in GSCs and differentiated GBM cells after treatment with TMZ as determined by extracellular flux analysis and XF Cell Mito Stress test Kit

(Seahorse, Agilent). A) Oxygen consumption rate (OCR) with kinetic graph and basal respiration, B) Proton leak, C) Maximal respiration, D) Spare respiratory capacity, and E) Non-mitochondrial oxygen consumption were determined. Values are shown as means ± SEM. Experiments were performed in two independent repeats (n = 2). Unpaired t-test was performed to compare statistical differences between control and treated samples within each cell line (A-F). * $p < 0.05$. Abbreviation: FCCP, Carbonyl cyanide 4-(trifluoromethoxy)phenylhydrazone.
